# Structure and iron-transporting mechanism of brain organic cation transporter 1

**DOI:** 10.1101/2025.06.28.662014

**Authors:** Jianan Chen, Shuhao Zhang, Wan Wei, Long Teng, Lei Tao, Kaixuan Gao, Bingqin Li, Aled M Edwards, Hao Fan, Xiangyu Liu, Ligong Chen

## Abstract

Brain organic cation transporter 1 (BOCT1), also known as the solute carrier family 22 member 17 (SLC22A17) or the receptor for lipocalin-2 (LCN2), plays critical roles in health and disease. Its deficiency in mice results in early postnatal mortality and severe neurogenesis impairments. Despite its importance in physiology and pathophysiology, BOCT1’s structure and transport mechanism remain elusive. Here, we integrate cryo-electron microscopy (cryo-EM), functional assays, biochemical experiments, and molecular dynamics simulations to elucidate the structure, substrate recognition, and transport mechanism of mouse BOCT1 (mBOCT1). The high-resolution cryo-EM structure reveals a distinctive N-terminal domain with a unique folding pattern dominated by a transmembrane loop atop TM6 (TML6), diverging from both known structures of SLC22 transporters and AlphaFold predictions. Notably, mBOCT1 functions as a high-capacity, low-affinity iron transporter independent of LCN2 binding. Iron transport is facilitated by a substrate gating mechanism involving TML6. These findings establish a structural basis for BOCT1’s role as an independent iron transporter, enhancing our understanding of the transport mechanisms within major facilitator superfamily (MFS) transporters and providing new insights into brain iron homeostasis.

## Introduction

As the second largest family of membrane proteins after G protein-coupled receptors (GPCRs), solute carrier proteins (SLCs) are pivotal in numerous physiological processes, including nutrient transport, neural signaling, immune responses, and tissue homeostasis^1–3^. Brain organic cation transporter 1 (BOCT1), classified as SLC22A17 within the SLC22 family, is highly expressed in the brain^4,5^. Previous studies link BOCT1 to cell apoptosis, multiple cancers, and Alzheimer’s disease (AD) ^6–9^. Recent findings reveal that conditional knockout of mouse BOCT1 (mBOCT1) causes severe weight loss, neural stem cell dysregulation, and early postnatal mortality, underscoring its essential role in brain physiology (unpublished data).

BOCT1 is primarily recognized as the receptor for lipocalin-2 (LCN2), an inflammatory mediator implicated in various pathologies^4^. LCN2 facilitates iron transport by binding siderophores^4,10,11^. Devireddy et al. proposed that BOCT1 binds iron-loaded LCN2, enhancing cellular iron levels via endocytosis, while binding to iron-deficient LCN2 reduces intracellular iron through exocytosis, potentially triggering cell apoptosis^4^. BOCT1 also interacts with other proteins, including metallothionein (MT), transferrin, and albumin^7^.

Compared to its role as a receptor, the transport mechanism and physiological functions of BOCT1 remain largely unexplored. As an “orphan” transporter, BOCT1 does not transport the typical organic metabolites of the SLC22 family, and the N-terminus of BOCT1 differs from other SLC22 members, as it lacks the typical transmembrane helices 1 (TM1). Additionally, the over 100 amino acids preceding the classical TM2 significantly differ from other proteins within the family (Supplementary Fig. 1)^12^. Although the N-terminus has been suggested to be crucial for the transport function of SLC22 family, discrepancies exist. For instance, OCT1/SLC22A1 retains full transport activity despite losing its first two TMs^13,14^. Furthermore, structural predictions suggest BOCT1 may contain 9, 11, or 12 TMs, reflecting uncertainty about its architecture^12,15^.

To address these gaps, we determined the first 3D structure of mBOCT1 using single-particle cryo-EM and validated its role as a distinct iron transporter through protein liposome transport and cellular uptake assays. Integrating cryo-EM structure with molecular dynamics (MD) simulations and site-directed mutagenesis, we uncovered mBOCT1’s substrate recognition and transport mechanism. This structural and mechanistic characterization provides a framework for understanding intracellular iron homeostasis and opens avenues for therapeutic targeting of BOCT1 in neurological diseases associated with iron dysregulation.

## Results

### Overall structure of mBOCT1

The full-length mBOCT1, consisting of 520 amino acids and sharing 96.5% sequence identity with human BOCT1 (Supplementary Fig. 2), was overexpressed in HEK293F cells and purified for structural analysis. The mBOCT1 cryo-EM structure was resolved to 3.3 Å overall resolution by single-particle analysis (Fig. 1a; Extended Data Figs. 1a, b). The structure, solved with C2 symmetry, revealed an antiparallel dimer configuration comprising 22 TMs (Extended Data Fig. 1). Further cell surface staining assays with N- and C-terminal GFP-tagged mBOCT1 indicated that this dimerization likely arises as an artifact of purification, with mBOCT1 adopting a cohabiting form in detergent micelles to maintain a lower energy state. Consistent with previous reports^12^, the N-terminus faces the extracellular side, while the C-terminus faces the intracellular side (Extended Data Fig. 2a). Interestingly, over 90 N-terminal residues remain unresolved in the cryo-EM structure, aligning with NMR studies that classify the N-terminus as an intrinsically disordered region (IDR)^12^.

**Fig. 1.**
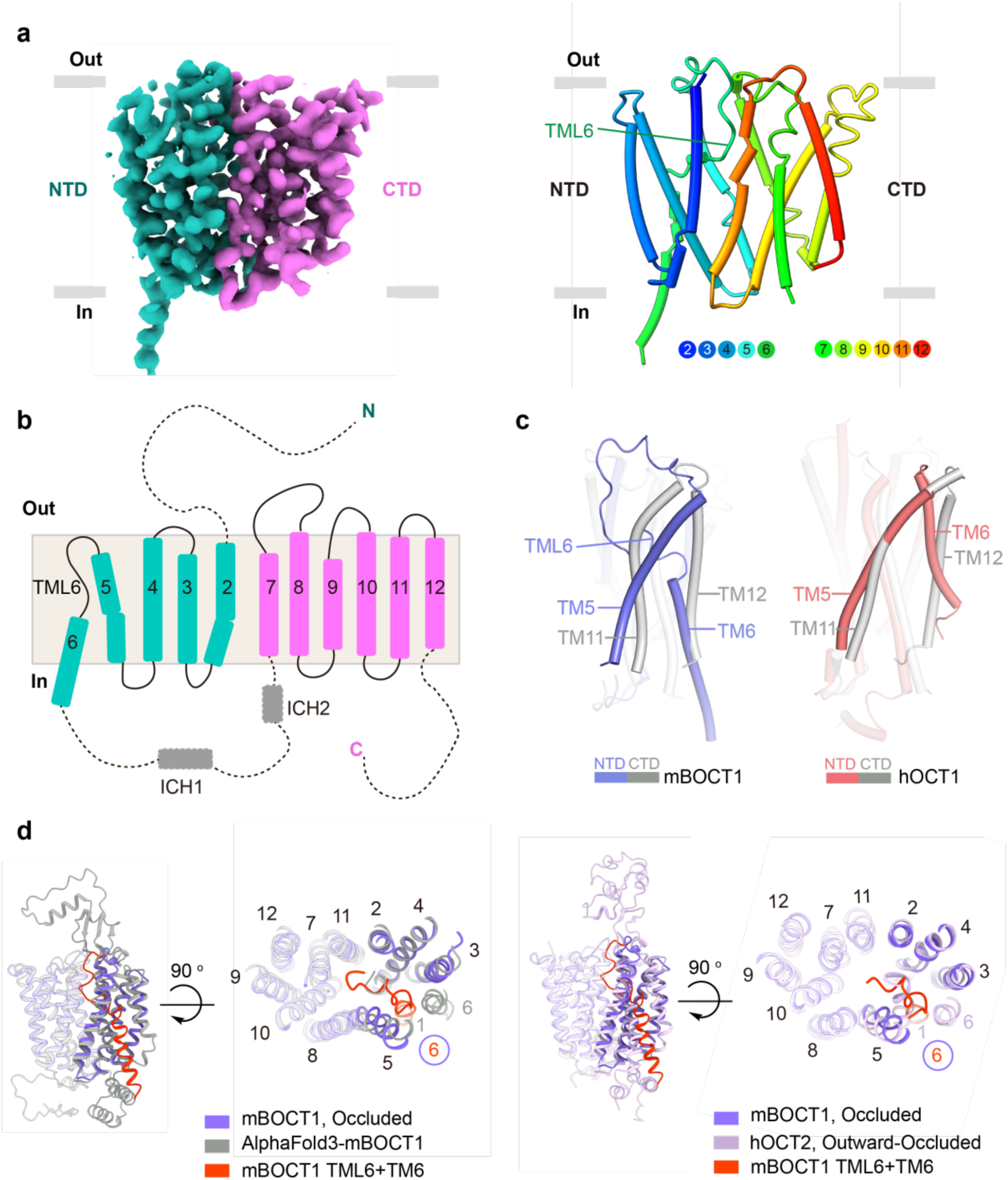
Overall structure of mBOCT1. **a**, Cryo-EM map and model of mBOCT1 monomer. Left: Cryo-EM map showing the NTD (green), CTD (magenta), and membrane environment (gray rectangles). Right: Model of mBOCT1 in rainbow colors, with each color corresponding to a TM (indicated by circles of matching color). TML6 is shown as a straight line. **b**, Topology of mBOCT1 monomer, with NTD (green) and CTD (magenta). The dashed line indicates unresolved regions. **c**, Structure alignments of NTD and CTD in mBOCT1 (this study) and hOCT1 (PDB: 8jtz). Non-transparent TMs (TM5, TM6, TML6, TM11, and TM12) are indicated by straight lines, while other regions are 80% transparent. mBOCT1 lacks pseudo-symmetric properties in its NTD and CTD. **d**, Structural alignment of mBOCT1 cryo-EM structure (purple) with AlphaFold3-predicted mBOCT1 (gray) and hOCT2 cryo-EM structure (PDB: 8et9) (purple). TML6 and TM6 in mBOCT1 are shown in red. Aligned TMs are labeled in black; unaligned segments are labeled in colors corresponding to their structures.

Transporters in the MFS typically possess 12 TMs with two-fold pseudosymmetry between their N-terminal domain (NTD; TM1-TM6) and C-terminal domain (CTD; TM7-TM12) ^16^. However, mBOCT1 comprises only 11 TMs, corresponding to TM2-TM12 of canonical MFS transporters. TM2-TM6 form the NTD, while TM7-TM12 constitute the CTD (Figs. 1a, b). Notably, the NTD and CTD of mBOCT1 lack the characteristic pseudosymmetry of MFS transporters due to distinct orientations of TM5-TM6 and TM11-TM12, which are usually aligned among other MFS members (Fig. 1c). Structural prediction corroborates the existence of a minority subset of MFS members with 11 TMs^16^, as evidenced by the crystal structure of human equilibrative nucleoside transporter 1 (hENT1) ^17^ that lacks TM12, while mBOCT1 lacks TM1.

A distinguishing feature of mBOCT1 is a disordered transmembrane loop (TML6; residues 220–236) located above TM6 in the cryo-EM structure, unlike homologous regions in experimental structures of other SLC22 members or AlphaFold3 predictions, which depict this segment as an intact helix distant from the transport pathway (Figs. 1b, c; Extended Data Fig. 2b). Nevertheless, sequence-based structural predictions suggest that TML6 might adopt a helical conformation under specific conditions (Supplementary Text in SI; Supplementary Fig. 3), and MD simulations reveal that TML6 can remain stable in both loop and helix forms (Extended Data Fig. 3a; Supplementary Table 4). Therefore, we speculate that mBOCT1’s TML6 may undergo conformational transitions between unique functional states (Extended Data Fig. 3a). The structural stability and helicity of all TMs of mBOCT1 in all-atom MD simulations show that TML6 is less stable than other TMs, and is more likely to undergo conformational rearrangements (Extended Data Fig. 3b). Moreover, TM6 had its helix elongated into the cytoplasm during certain segments of MD simulations (Extended Data Fig. 3c). Collectively, we speculate that substrate transport of mBOCT1 is accompanied by conformational changes of both TML6 and TM6.

### Substrate Identification of mBOCT1

Electrostatic analysis reveals that the transport cavity of mBOCT1 is negatively charged, similar to the organic cation transporter OCT1^18^ and the Fe^2+^ efflux transporter hFPN1^19^, but distinct from the organic anion transporter OAT1 (Fig. 2a; Extended Data Figs. 4a, b)^20^. This suggests that mBOCT1 likely transports positively charged substrates. The binding pocket of mBOCT1 also exhibits hydrophilic characteristics similar to that of hFPN1^19^, but differs from those of OCT1 and OAT1 (Extended Data Figs. 4a, c)^18,20^. Conservation analysis performed using Consurf highlights significant differences in mBOCT1’s binding pocket compared to other SLC22 family members (Fig. 2b).

**Fig. 2.**
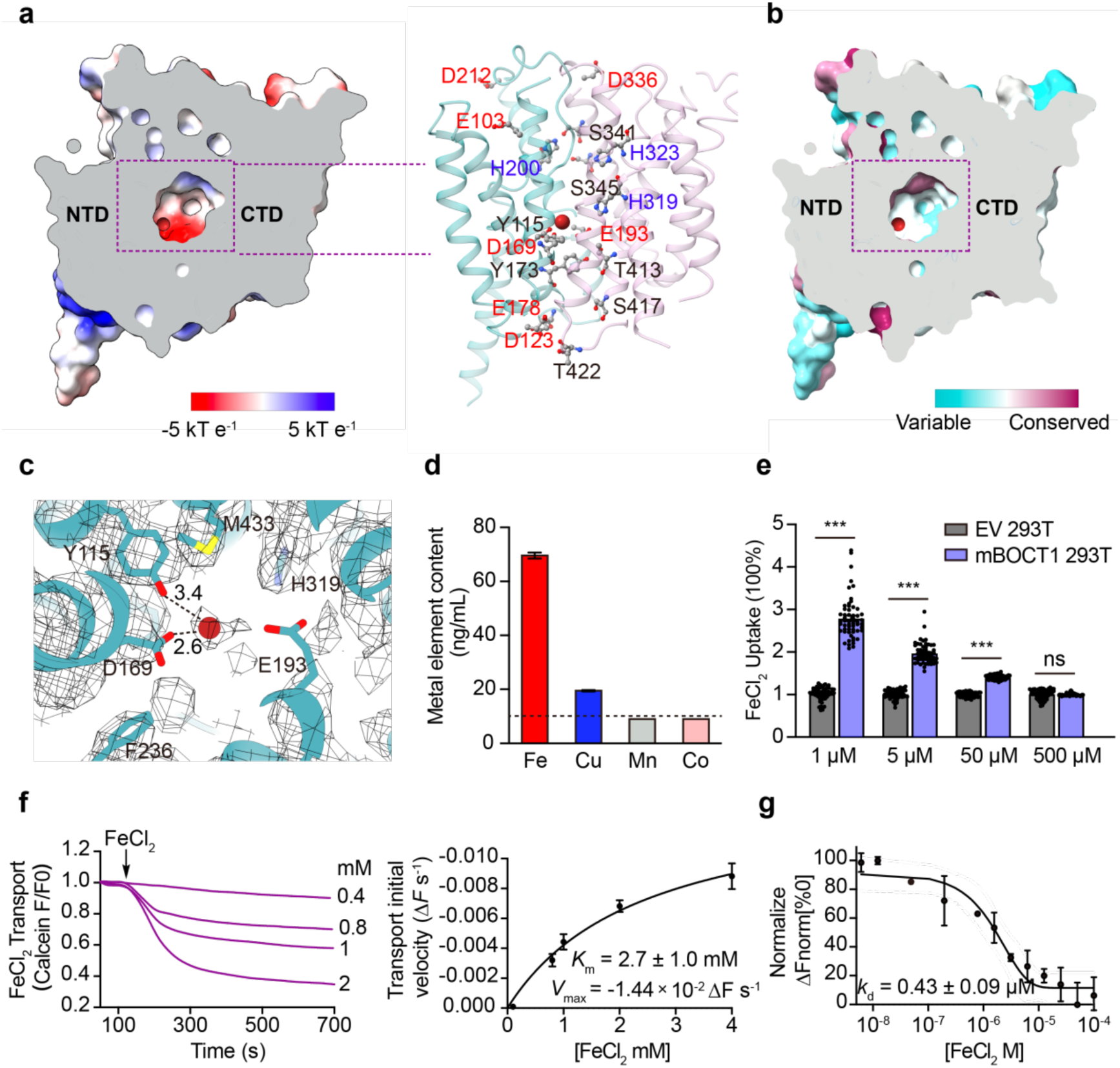
Structural analysis and substrate identification of mBOCT1. **a**, Sliced view of the binding pocket showing surface electrostatic potential and polar residues. Fe^2+^ ion is shown as a red sphere; amino acids are depicted as sticks. The color bar indicates electrostatic potential. **b**, Sliced view showing variability of mBOCT1 compared to selected SLC22 family members, calculated by Consurf. The Fe^2+^ ion is shown as a red sphere; the color bar represents conservation levels. **c**, Extra density and surrounding residues in the mBOCT1 transport pathway. **d**, Metal ion content in mBOCT1 (cryo-EM sample preparation), detected by ICP-MS. Metal content below 10 ng/mL (dashed line) is considered unreliable. Data are presented as mean ± SD (n = 3). **e**, Cellular FeCl_2_ uptake assay in mBOCT1-expressing HEK293T cells, normalized to EV controls. Data are shown as mean ± SEM (n>3). Unpaired *t*-test: \**p* < 0.05, \*\**p* < 0.01, \*\*\**p* < 0.001. **f**, Fe^2+^ transport by mBOCT1 in calcein-containing liposomes. Time-course curves are averaged from n=3 independent experiments. Initial velocity was obtained from the linear phase of the time-course curve and fitted using the Michaelis-Menten formula in GraphPad Prism. Data are Mean ± SD. **g**, FeCl_2_ binding affinity of mBOCT1, detected by MST. Data are mean ± SD (n = 3).

We observe an additional density within the binding pocket, and the binding pocket comprises a mix of polar and aromatic residues suitable for metal ion coordination, including Y115, D169, and E193 in close proximity and H319 slightly farther away (Figs. 2a, c). Using inductively coupled plasma mass spectrometry (ICP-MS), we identified iron as the predominant metal ion associated with mBOCT1 protein samples analyzed via cryo-EM (Fig. 2d).

To validate its metal ion transport activity, we conducted a cellular uptake assay using calcein quenching. mBOCT1-expressing HEK293T cells exhibited 2–3 times greater FeCl_2_ transport activity compared to empty vector (EV) controls, with similar results for FeSO_4_ (Fig. 2e; Extended Data Fig. 5). Protein-liposome transport assays further confirmed mBOCT1’s ability to transport FeCl_2_ with a Michaelis-Menten constant (*K*_m_) of 2.7 ± 1.0 mM and *V*_max_ of −1.44×10^−2^ ΔF s^−1^ (Fig. 2f). mBOCT1 liposomes also transported other metal ions, including Mn^2+^ and Co^2+^ (Extended Data Figs. 6a-c). In addition, Microscale thermophoresis (MST) assays that measure binding affinities, revealed a dissociation constant (*k*_d_) of 0.43 µM for FeCl_2_, comparable to CuCl_2_ and MnCl_2_, while CoCl_2_ exhibited a higher binding affinity (Fig. 2g; Extended Data Fig. 6d). These findings establish mBOCT1 as a transporter for multiple divalent metal ions, facilitating their cellular uptake independently.

### Molecular basis of mBOCT1 substrate recognition

To confirm the iron-binding site identified in the mBOCT1 cryo-EM structure, we introduced mutations in the putative binding pocket and measured their FeCl₂ transport activity. Mutations D169A, D169T, and E193A resulted in substantially reduced activity (~20% of wildtype [WT]), highlighting the critical role of the negative charges on D169 and E193 in facilitating iron transport. Y115 likely interacts with iron through polar and cation-π interactions, as the Y115F mutant retained ~40% of WT activity (Fig. 3a). MD simulations supported this, showing that Fe^2+^ remained stable at this site (Figs. 3b, c; Supplementary Table 5) and frequently interacted with the binding site residues (Supplementary Table 6).

**Fig. 3.**
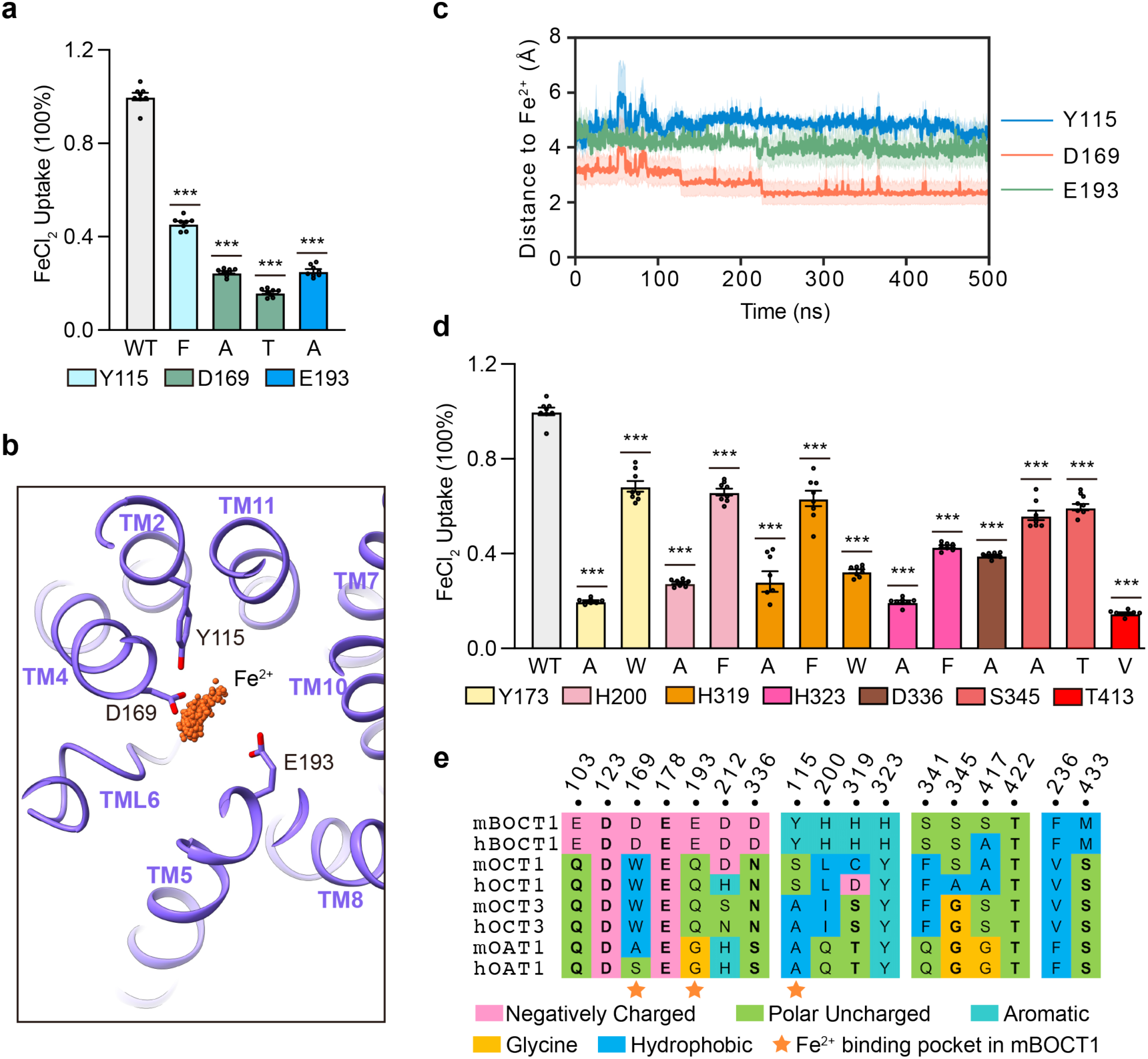
Substrate binding verification in mBOCT1 through mutations and MD simulations. **a**, Fe^2+^ transport activity of binding pocket mutations. Data are mean ± SEM (n>3). Unpaired *t*-test: \**p* < 0.05, \*\**p* < 0.01, \*\*\**p* < 0.001. **b**, Fe^2+^ in the mBOCT1 binding pocket from all-atom MD simulations. **c**, Distances between Fe^2+^ and binding site residues (Y115, D169, E193) during 5 replicas of 500 ns holo MD simulations. Solid lines show the mean; shaded bands indicate SE across 5 replicas. **d**, Fe^2+^ transport activity of pathway mutations. Data are mean ± SEM (n>3). Unpaired *t*-test: \**p* < 0.05, \*\**p* < 0.01, \*\*\**p* < 0.001. **e**, Sequence alignment of key Fe^2+^-transport residues between mBOCT1 and selected SLC22 members. Residues are color-coded by properties.

Mutating polar residues along the transport pathway (e.g., Y173A, H200A, H323A, T413V, Supplementary Fig. 4) significantly impaired FeCl₂ transport, underscoring their pivotal roles. H319A displayed impaired activity similar to D169A and E193A, while H319F retained ~60% of WT activity, suggesting that H319 contributes through polar and cation-π interactions. However, the H319W mutation caused significant activity loss, likely due to steric hindrance. Similarly, H200F and H323F mutations had less severe reductions in activity than their alanine counterparts, possibly due to favorable cation-π interactions with aromatic rings (Fig. 3d). These key residues are conserved in human and mouse BOCT1 but not in other SLC22 family members, indicating functional conservation across species (Fig. 3e).

### Iron transport gating mechanism of mBOCT1

In the mBOCT1 cryo-EM structure, the negatively charged residue D336 is positioned at the entrance of the proposed transport pathway, formed by the NTD and CTD (Fig. 2a). The D336A mutation reduced FeCl₂ transport activity to ~40% of WT levels (Fig. 3d), suggesting that D336 acts as a “hook” to attract iron, a hypothesis supported by MD simulations (Extended Data Fig. 7a, Supplementary Movie 1).

mBOCT1’s TML6 is situated at the entrance of the transport pathway, differing from the mBOCT1 AlphaFold3 model or experimental structures of other SLC22 family members (Fig. 1)^21^. mBOCT1 features a unique GGGG (329-332) motif, which is rare within the SLC22 family (Extended Data Fig. 2b). These four consecutive glycine residues are situated near TML6 along the transport pathway. The TML6 and GGGG motif constitute an extracellular gate for mBOCT1 (Figs. 4a, b). In MD simulations based on the cryo-EM structure, Fe^2+^ was unable to penetrate the pathway unless TML6 moved away to allow substrate entry, emphasizing TML6’s role in gating (Figs. 4a, c; Supplementary Fig. 5). TML6 is stabilized by hydrophobic interactions between F229 (TML6) and F107/F111 (TM2), that possibly contributes to the transporter’s occluded conformation (Figs. 4a, d). On the intracellular side, Y173 and T413 form sidechain hydrogen bonds and may act as a gate, controlling Fe^2+^ release into the cell (Figs. 4a, e).

**Fig. 4.**
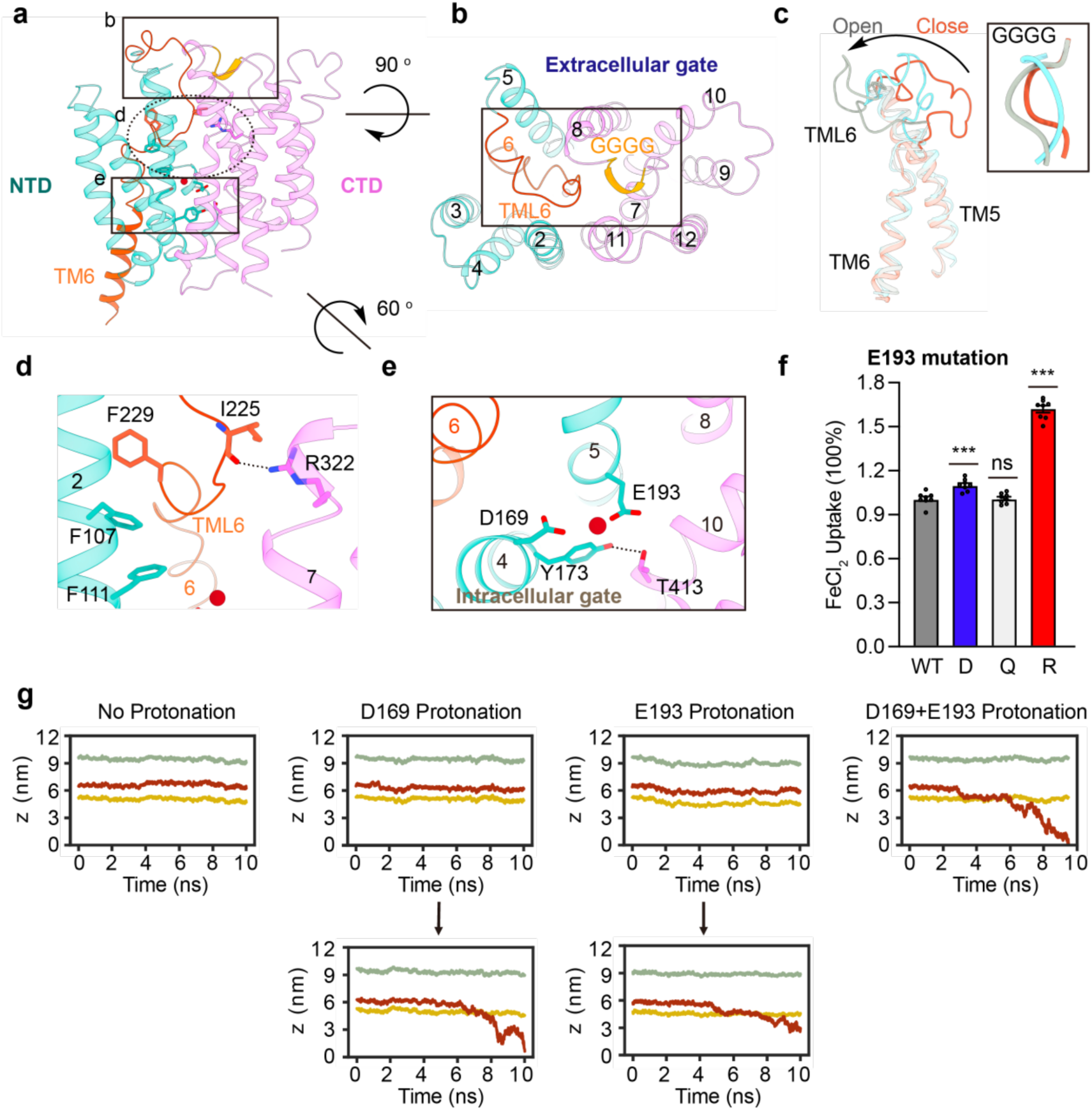
Substrate transport mechanism of mBOCT1. **a**, Side view of the overall mBOCT1 structure. **b**, Top view of mBOCT1; the black box highlights the b region shown in panel a. TML6 and the GGGG motif form the extracellular transport gate. **c**, Structural diversity of TML6 in coarse-grained MD simulations. TML6 in the open state (gray) permits Fe^2+^ entry; in the closed state (red), it blocks transport. **d**, Zoomed view of region d (panel a) showing TML6 interactions with other TMs. Hydrogen bond: I225 (TML6) with R322 (TM7). Hydrophobic interactions: F229 (TML6) with F107 and F111 (TM2). **e**, Zoomed view of region e (panel a) showing the intracellular transport gate formed by Y173–T413 hydrogen bonding. **f**, Fe^2+^ transport activity of E193 mutations. Data are mean ± SEM (n>3). Unpaired *t*-test: \**p* < 0.05, \*\**p* < 0.01, \*\*\**p* < 0.001. **g**, Protonation effects on Fe^2+^ transport in mBOCT1 observed in parallel cascade selection molecular dynamics (PaCS-MD) simulations. PaCS-MD was conducted to investigate the Fe^2+^ ion transport in mBOCT1 through multiple rounds of short parallel simulations. For each round, 100 short MD simulations of 10 ns each were conducted. All MD frames were collected and ranked by the penetration depth of Fe^2+^ ions in the lipid bilayer. The top-ranked frame was utilized as the starting frame for the next round of PaCS-MD simulations. For each system (System 11.1-11.4 in Supplementary Table 2), the Fe^2+^ trajectory from the simulation with the deepest ion penetration is presented in the plot. Green: extracellular phospholipid layer; yellow: intracellular layer; red: Fe²⁺ trajectory.

### Protonation on iron transport in mBOCT1

Mutating E193 to aspartate or asparagine did not affect transport activity, while replacing it with arginine enhanced activity (Fig. 4f). This indicates that the protonation state of E193 affects Fe^2+^ release, as supported by parallel cascade selection molecular dynamics (PaCS-MD) simulations (Supplementary Table 2). In these simulations, protonation of D169 or E193 increased the distance between these two residues, allowing greater Fe^2+^ mobility within the transport pathway (Fig. 4g), as indicated by the altered angle formed by Fe^2+^, D169, and E193 (Extended Data Fig. 7b). The average and maximum Fe^2+^ ion depth in the lipid bilayer also increased (Supplementary Table 7). Thus, the Fe^2+^ ion is more likely to be released from the binding pocket when D169 or E193 is protonated.

In addition, MD simulations revealed that Fe^2+^ is consistently surrounded by a hydration shell of 5-6 water molecules (Extended Data Figs. 7c, d), which forms hydrogen bonds with D169 and E193. Protonation of these two residues would disrupt these hydrogen bonds and facilitate Fe^2+^ release, with a more pronounced effect upon E193 protonation, which affected water hydrogen bonding with both D169 and E193. In contrast, D169 protonation disrupted only its own hydrogen bonds with water (Extended Data Figs. 7d, e). A recent study has demonstrated that the protonation and deprotonation of D/E residues regulate extracellular gating in a homologous organic anion transporter, SLC17^22^, supporting our hypothesis that D/E residue protonation influences mBOCT1 iron transport. However, further studies are needed to fully elucidate this mechanism.

### Fe^2+^ transport model of mBOCT1

Structural comparisons between the “outward-open” hOCT1^21^, “inward-open” hOCT1^23^, and mBOCT1 reveal that TML6 in mBOCT1 aligns spatially with TM1, rather than TM6 of classic MFS transporters (Extended Data Figs. 8a, b). Given the established importance of conformational changes in TM1 for substrate transport in classic MFS transporters^16^, this highlights the critical role of TML6 in mBOCT1 substrate transport. Additionally, conformational changes in TM7 and TM10 of mBOCT1 are likely integral to the transport process (Extended Data Figs. 8a, b). The aforementioned changes were also observed when we conducted structural alignments between the homology modeling-generated outward-open mBOCT1 model, inward-open mBOCT1 model, and the cryo-EM structure of the occluded mBOCT1 (Supplementary Fig. 6; Supplementary Text in SI).

Building on these observations and the rocker-switch substrate transport mechanism of MFS transporters, we propose a new Fe^2+^ transport model for mBOCT1. Initially, in the hypothesized outward-open conformation, TML6 and the GGGG motif orient away from the transport pathway, leaving the extracellular gate open. Fe^2+^, attracted by electrostatic interactions with negatively charged residues such as D336, enters the pathway. Subsequently, interactions between TML6 and other TMs shift TML6 toward the transport pathway, preventing further substrate entry. The extracellular gate is sealed by a hydrogen bond between I225 on TML6 and R322 on TM7.

Once inside, Fe^2+^ is progressively guided downward by polar residues such as H200 and H319, followed by negatively charged residues D169 and E193. Protonation of D169 or E193 likely triggers conformational changes in TM7 and TM10 of the CTD, potentially coupled with TM6 extending into the cytoplasm. This facilitates the opening of the intracellular gate, formed by hydrogen bonds involving Y173 and T413, allowing Fe^2+^ release into the cell and completing the transport cycle (Fig. 5).

**Fig. 5.**
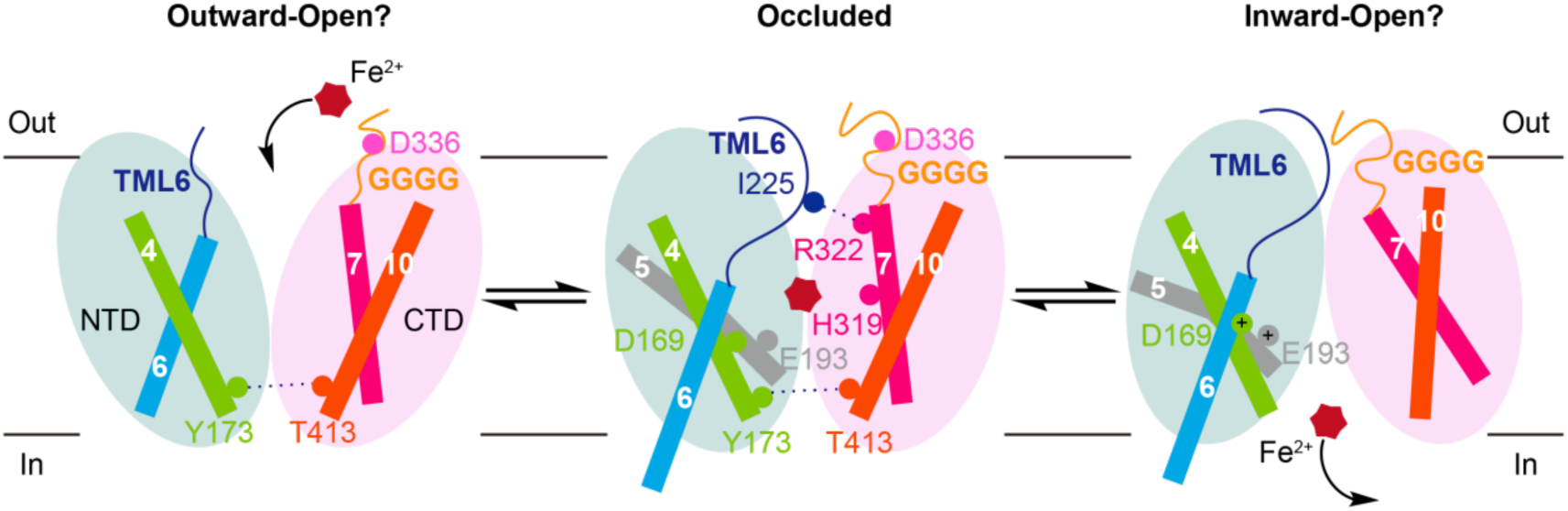
Hypothetical mBOCT1 Fe^2+^ transport model. The outward-open and inward-open conformations of mBOCT1 are hypothetical, and the occluded conformation is a simplified representation of the mBOCT1 cryo-EM structure. The NTD and CTD of mBOCT1 are shown in light green and light red, respectively. TMs are represented as rectangles in distinct colors, loops as lines, amino acid residues as spheres, and Fe^2+^ as a hexagon. Plus signs indicate protonation, while “out” and “in” denote the extracellular and intracellular environments, respectively.

## Discussion

This study presents the high-resolution cryo-EM structure of mBOCT1 in complex with endogenous Fe^2+^ in its occluded conformation. Unlike typical MFS transporters, which exhibit pronounced pseudosymmetry between their NTD and CTD^16^, mBOCT1’s NTD and CTD adopt folding patterns that diverge significantly from other MFS members.

Notably, mBOCT1 demonstrates the ability to transport Fe^2+^ independently of LCN2. Previous studies suggest that LCN2 binds to siderophores and facilitates Fe^3+^ transport via BOCT1^4^. However, our findings indicate that mBOCT1 selectively transports divalent metal ions such as Fe^2+^, highlighting its dual functionality: receptor-mediated Fe^3+^ transport and transporter-mediated Fe^2+^ transport. The substrate affinity (*K_m_)* for Fe^2+^, determined via liposome transport assays, is approximately 2.7 mM—considerably lower than the *K_m_* of 13.6 µM reported for the classical Fe^2+^ efflux transporter hFPN1 (SLC40A1)^19^. Conversely, mBOCT1’s Fe^2+^ binding dissociation constant (*k_d_*), measured by MST, is ~0.43 µM, significantly stronger than the 30 µM *k_d_* observed in the bivalent metal transporter ScaDMT (SLC11 family)^24^. These results classify mBOCT1 as a “High Capacity, Low Affinity” transporter, akin to the OCT family, in contrast to the “High Affinity, Low Capacity” transporters of the SLC6 family^25,26^.

Unlike other SLC22 family members, which primarily recognize substrates through aromatic residues such as Y230 and F438 in OAT1 or Y36/Y37 and Y361/Y362 in OCT1-2^20,21^, mBOCT1 engages residues Y115, D169, and E193 for Fe^2+^ recognition. Additionally, while TM7, TM10, TM11, and intracellular loop (ICH) regions in OCTs and the aromatic residues Y353 and F442 in OAT1 facilitate substrate transport through conformational changes^20,23^, our data suggest that mBOCT1 employs a unique mechanism where TML6 serves as a pivotal gating element (Fig. 4, 5).

Sequence alignment further supports that hBOCT1 likely mirrors mBOCT1’s transport mechanism, given the high conservation of key residues involved in substrate recognition (Y115, D169, E193), TML6 folding (F107, F111, F229), and the formation of the GGGG motif. In contrast, corresponding homologous residues in human and mouse OCT1, OCT3, and OAT1 show limited conservation, with the exception of F111 (Fig. 3e; Extended Data Fig. 2b). This divergence underpins the functional differences between BOCT1 and other SLC22 family members.

In summary, this study identifies Fe^2+^ as the direct substrate of BOCT1, delineates key residues involved in substrate recognition and transport, and proposes a distinct molecular transport mechanism. Our findings advance the “de-orphaning” of the SLC transporter family and highlight BOCT1 as a potential therapeutic target for iron metabolism and homeostasis-related disorders. Dysregulation of iron homeostasis is closely linked to neurodegenerative diseases such as multiple sclerosis, Parkinson’s disease, and Alzheimer’s disease^27–29^. The structural and functional insights provided here deepen our understanding of these pathologies and may guide rational drug design. Finally, while AlphaFold^30,31^ has achieved remarkable accuracy in structure prediction, the unexpected cryo-EM structure of BOCT1 underscores the indispensable role of traditional structural biology methods.

**Table 1.** Cryo-EM data collection, refinement, and validation statistics.

|  | *m*BOCT1 |
| --- | --- |
| **Data collection and processing** |  |
| Microscope | Titan Krios G3i |
| Camera | K3 |
| Spherical Aberration (mm) | 0.01 |
| Magnification | 64000 |
| Voltage (kV) | 300 |
| Electron exposure (e-/Å<sup>2</sup>) | 50 |
| Exposure time (s) | 2.53 |
| Defocus range (μm) | -1.2 to -1.5 |
| Pixel size (super resolution Å) | 0.54 |
| Slit width (eV) | 20 |
| Frame | 32 |
| Micrographs | 4649 |
| Autopicked particles | 7659501 |
| Particles in final refinement | 178900 |
| Symmetry imposed | C2 |
| FSC threshold | 0.143 |
| Map sharpening B factor (Å<sup>2</sup>) | -138.7 |
| Map resolution (Å) | 3.3 (cryoSparc) |
| Map resolution range (Å) |  |
| FSC 0.5, unmasked/masked | 6.3-3.9 |
| FSC 0.143, unmasked/masked | 3.5-3.4 |
| **Refinement** |  |
| Initial model used (PDB code) | - |
| Model composition |  |
| Non-hydrogen atoms | 4946 |
| Protein residues | 666 |
| Ligands | Fe:2 |
| Water | N/A |
| *B* factors (Å<sup>2</sup>) |  |
| Protein | 99.54 |
| Ligand | 93.71 |
| Water | - |
| R.m.s. deviations |  |
| Bond lengths (Å) | 0.003 |
| Bond angles (°) | 0.622 |
| **Validation** |  |
| Rotamers outliers (%) | 0 |
| Molprobity score | 2.05 |
| CaBLAM Score | 2.02 |
| Clashscore | 13.08 |
| Ramachandran plot |  |
| Favored (%) | 93.62% |
| Allowed (%) | 6.38% |
| Disallowed (%) | 0 |

## Methods

### Protein expression and purification of mBOCT1

The mBOCT1 protein (NCBI: NP_001347335.1), tagged at the C-terminus with either a strep tag or GFP and a 3C protease cleavage site, was cloned into the pcDNA3.1(+) vector. The recombinant plasmid, free from endotoxins, was transfected into HEK293F cells using PEI at a 3:1 mass ratio. After 48 hours, cells were harvested and lysed using a buffer containing 20 mM Tris-HCl pH 8.0, 2 mM EDTA, and protease inhibitors. The lysate was centrifuged, and the resulting pellet was solubilized in 20 mM Hepes pH 8.0, 100 mM NaCl, 1% (w/v) MNG, 0.1% (w/v) CHS, and protease inhibitors before further centrifugation at 18,000 rpm.

The supernatant was applied to strep affinity beads and homemade anti-GFP nanobody Sepharose beads, equilibrated and rinsed in 20 mM Hepes pH 8.0, 100 mM NaCl, 0.01% (w/v) MNG, and 0.001% (w/v) CHS. For strep-tagged mBOCT1, the protein was eluted with the same buffer supplemented with 2.5 mM Desthiobiotin. For GFP-tagged mBOCT1, the tag was removed using 3C protease. The eluted protein was concentrated with a 50-kDa MWCO Millipore concentrator and subjected to size-exclusion chromatography (SEC) using a Superdex 200 Increase column (GE) equilibrated in 20 mM Hepes pH 8.0, 100 mM NaCl, 1 mM TCEP, 0.003% (w/v) MNG, and 0.0003% (w/v) CHS. Peak fractions were collected for experiments, and purity was confirmed by SDS-PAGE.

### Cryo-EM sample preparation and data collection

Purified mBOCT1 peak fractions were concentrated to 5 mg mL⁻¹. Four µL of the sample was applied to glow-discharged Quantifoil gold 200 mesh grids (R1.2/1.3), blotted for 4.5 s and waited for 5 s, and plunge-frozen in liquid ethane using a FEI Mark IV Vitrobot at 8 °C and 100% humidity. Data collection was performed on a 300 kV Titan Krios G3i equipped with a BioQuantum K3 camera. Movies were recorded at 50 e⁻/Å^2^ with a defocus range of −1.2 to −1.5 µm. A total of 4,649 micrographs were collected with a magnification of 64,000x, corresponding to a pixel size of 1.08 Å.

### Cryo-EM data processing

Data were processed using cryoSPARC 3.2 and 4.1^32^. An initial 185 micrographs underwent patch CTF estimation, and micrographs with estimated CTF resolutions worse than 4 Å were excluded after checked by manual curate exposure. After two rounds of 2D classification, selected classes were used for template picking in the entire set of 4649 micrographs, yielding 7,433,396 particles extracted at 256 pixels and downsampled to 128 pixels. Ab initio models were generated from good classes or bad classes, followed by heterogeneous refinement to remove suboptimal particles. The final dataset consisted of 360,940 particles re-extracted at 256 pixels, then the 2D classification and heterogenous refinement were repeated, and 171,512 particles were used for final refinement. Local CTF refinement was applied to per-particle. Finally, a 3.3 Å C2 symmetry density map was obtained.

### Model building and refinement

The initial mBOCT1 model was generated using cyronet.ai and AlphaFold, followed by iterative refinement in Coot and real-space refinement in PHENIX^33,34^. Validation was performed using PHENIX and MolProbity, with figures generated in PyMOL and ChimeraX^35^.

### Identification of metal element content in purified mBOCT1 protein

The metal content of purified mBOCT1 was analyzed via ICP-MS (Agilent Technologies). Equal volumes of 1.2 mg mL^−1^ mBOCT1 and control buffer were mixed with 100 µL of high-purity nitric acid and heated for 4 hours in a water bath. The reaction mixtures were adjusted to a final volume of 5 mL and analyzed via ICP-MS.

### Surface staining by immunofluorescence

HEK293T cells were stably transfected with mBOCT1 constructs tagged with GFP at the N-terminus or C-terminus or an empty vector (EV). Cells were seeded on poly-D-lysine-coated slides and blocked with 1% BSA at room temperature for 30 minutes. Cells were stained with 100 nM Alexa647-GFP nanobody, washed with PBS, and fixed with 4% PFA. Nuclei were counterstained with 1 µg µL^−1^ DAPI, and slides were sealed with an anti-fluorescent quencher agent. Samples were imaged using an Olympus FV3000 confocal microscope.

### Construction of mBOCT1 stable cell line

The wild-type (WT) and mutant mBOCT1 constructs, each bearing an N-terminal HA and Flag tag, were cloned into the lentiviral vector pLJM1. The recombinant pLJM1 plasmid, along with the packaging plasmids pMD2.G and psPAX2 (mass ratio 5:3:2), was co-transfected into HEK293T cells using PEI. Lentiviral supernatants from these cells were harvested and used to infect fresh HEK293T and CHO cells. Puromycin (Beyotime) was added at appropriate concentrations to select for positively transfected cells. After a two-week screening process, surviving cells were deemed successfully transfected. For GFP-tagged mBOCT1 constructs, an additional step of positive cell sorting was performed using a FACSAriaII (BD). Overexpression efficiency was confirmed via qPCR and western blot.

### Cellular substrate uptake assay

Stable HEK293T and CHO cell lines expressing mBOCT1 with N-terminal HA and Flag tags, as well as cells transfected with the empty vector (pLJM1), were seeded into 96-well plates. On the day of the assay, cells were washed with PBS, followed by the addition of 100 µL calcein-AM (1 µM, diluted in serum-free medium) and incubation at 37 °C for 40 minutes. The dye solution was removed, and the cells were incubated in fresh serum-free medium at 37 °C for another 40 minutes to ensure the complete conversion of non-fluorescent calcein-AM to fluorescent calcein. After washing with HBSS (without calcium and magnesium), baseline fluorescence was recorded using a microplate reader (excitation: 490 nm, emission: 520 nm) for at least 1 minute. Subsequently, 10 µL of a solution containing metal ions diluted in HBSS (without calcium and magnesium) was added, and the uptake curve was measured over 4 minutes.

For dose-response experiments, FeCl_2_ concentrations ranged from 1 µM to 500 µM. For iron transport assays with mBOCT1 mutants, Fe^2+^ and other metal ions were used at final concentrations of 2 µM. Sodium ascorbate was added at a 20-fold molar ratio to ferric ions to maintain a reductive environment. Each experiment was independently repeated in triplicate, and data were analyzed using GraphPad Prism.

### Protein liposome assembly

Protein liposome preparation followed the method by Billesbølle et al ^19^. Lipids POPE and POPG were dissolved in chloroform (mass ratio 3:1), dried under argon gas, resuspended in an internal buffer (20 mM HEPES, pH 7.4, and 100 mM KCl), and sonicated. The lipid mixture underwent repeated freeze-thaw cycles and extrusion through a 400 nm polycarbonate filter. To destabilize liposomes, 0.13% (w/v) Triton-X-100 was added, followed by purified mBOCT1 (lipid-to-protein ratio of 40:1). After incubation at 4 °C for 30 minutes, SM2-Biobeads (0.05 mg/mL) were used to remove detergent over a 1-hour period. This process was repeated overnight, followed by a final incubation with 0.08 mg/mL SM2-Biobeads for 2 hours at 4 °C. Protein liposomes were collected by centrifugation at 246,000 × *g* for 30 minutes, resuspended in internal buffer, and frozen by liquid nitrogen, and then stored at −80 °C. A control sample, prepared without purified mBOCT1, was similarly processed.

### Protein liposome-based transport assay

On the experiment day, liposomes were thawed and resuspended at 2 mg/mL in an internal buffer containing 250 µM calcein. Liposomes were subjected to repeated freeze-thaw cycles, extrusion, and concentration at 264,000 × *g* for 30 minutes. Calcein-containing liposomes were washed with an external buffer (20 mM HEPES, pH 7.4, and 100 mM NaCl).

During the assay, 100 nM valinomycin was added to establish a membrane potential. Liposomes were added to a final concentration of 0.1 mg/mL. For dose-response experiments, FeCl_2_ concentrations ranged from 0.05 µM to 5 mM. Other metal ions were tested at 1 mM. Sodium ascorbate (20-fold molar ratio) was included to maintain a reductive environment for ferric ions. Fluorescence was recorded at 2–3 s intervals (excitation: 490 nm, emission: 520 nm) over at least 1 minute. Transport was terminated by adding 10 µM calcimycin, and data were collected using a microplate reader. All experiments were independently repeated in triplicate, and results were analyzed with GraphPad Prism.

### Binding assay of mBOCT1 with substrates

The binding interaction between mBOCT1 and metal ions was analyzed using MST. Purified mBOCT1 was labeled with a Monolith™ RED-NHS secondary generation protein labeling kit, and unbound dye was removed following the manufacturer’s protocol. Prior to labeling, mBOCT1 underwent buffer exchange to minimize TCEP interference, using a replacement buffer containing 20 mM Hepes pH 8.0, 100 mM NaCl, 0.003% (w/v) MNG, and 0.0003% (w/v) CHS.

For the assay, 10 µM mBOCT1 was labeled, and 40 nM of the labeled protein was mixed with serial dilutions of CoCl_2_ (25 µM to 1.5 nM) and other metal ions (100 µM to 3.1 nM). To ensure a reductive environment for ferric ions, sodium ascorbate was added at a 20-fold molar ratio. The mixtures were centrifuged at 13,000 rpm and loaded into Monolith™ NT115 capillaries (K-022). The MST experiments were conducted under medium power and 60% auto-detection using the Monolith™ NT115 system from NanoTemper Technologies. Data were analyzed with Mo.Affinity Analysis software (X86), and each experiment was independently repeated three times.

### Modeling of TML6 and unresolved intracellular domain by MD simulations

The experimentally determined structure was refined using the Bayesian inference-based EMMIVox approach implemented in PLUMED (www.plumed.org) ^36,37^. Missing atoms were reconstructed using the CHARMM-GUI web server^38,39^, which also generated topology files. The cryo-EM map was aligned with the reconstructed conformation in PLUMED format.

Structural refinement involved minimization and equilibration (1 ns NPT, followed by 2 ns NVT) at 300 K, with position restraints of 400 kJ/(mol·nm^2^) on the protein backbone and 40 kJ/(mol·nm^2^) on sidechain heavy atoms. The NVT equilibration trajectory was used to optimize the scaling factor between the experimental cryo-EM map and the derived map. A 10 ns NVT production run followed, applying restraints from the cryo-EM map. The conformation with the lowest EMMIVox hybrid energy was extracted along with its B-factors. This conformation underwent steepest descent energy minimization, sampling B-factors via a modified Monte Carlo approach. The final minimized structure was re-aligned to the original cryo-EM map, with sampled B-factors incorporated to generate a complete PDB file containing heavy atoms only. This refined cryo-EM model was used for following modeling and simulations. All simulations were performed using GROMACS 2021.5, integrated with PLUMED^40^.

To model the unresolved intracellular domain (residues 258–302), RoseTTAFold on the Robetta server (https://robetta.bakerlab.org/) was employed for structure prediction^41^. Based on secondary structure and transmembrane region predictions from the protein sequence^42^, TML6 (residues 220-236) was predicted to be a transmembrane helix, as opposed to the loop observed in the cryo-EM map (Supplementary Fig. 3). As such, two models were used to represent the mBOCT1 protein (residues 98-475). In the first model (model A), the intracellular domain structure was taken from the Robetta model and added to the refined cryo-EM structure. In the second model (model B), TML6 was presented as a helix, and both the TML6 helix and the intracellular domain were combined with other transmembrane helices from the refined cryo-EM structure. MODELLER was used to combine the Robetta predictions with the refined cryo-EM structure^43^. For each model, 5000 models were generated and refined, while the one with the best DOPE score was selected for further optimization using Maestro’s protein preparation wizard (Schrödinger, NY). The optimized model A and B were used for subsequent simulations.

### All-atom molecular dynamics (MD) simulations

All-atom MD simulations were performed on the refined cryo-EM structure of mBOCT1, incorporating the intracellular domain predicted by the Robetta model. These simulations were carried out in explicit solvents and lipid bilayers composed of 70% POPC and 30% cholesterol by molar ratio^44^. The protein was first aligned within the membrane using the PPM 2.0 Web server before system assembly in CHARMM-GUI^38,45–47^. The system was solvated in water, with Na⁺ and Cl⁻ ions added to neutralize the charge and maintain a physiological NaCl concentration of 150 mM. For the iron-bound (holo) system, one Fe^2+^ ion was added along with balancing Cl⁻ ions.

The CHARMM36 force field was used to describe the protein, lipids, and ions, while water was modeled using the CHARMM TIP3P water model^48,49^. Simulations were conducted both with and without Fe^2+^ bound to the protein, and all simulation systems are detailed in Supplementary Tables 1 and 3. After equilibration following the CHARMM-GUI Membrane Builder protocol at 310.15 K^46,47^, 3 replicas of 1 µs and 15 replicas of 200 ns production runs were performed for apo systems, while 5 replicas of 500 ns runs were conducted for holo systems to assess ion binding. All productive runs were performed in the NPT ensemble, maintaining a temperature of 310.15 K via Nosé-Hoover thermostat and a pressure of 1 bar using Parrinello-Rahman coupling. Lennard-Jones interactions were truncated between 10 Å and 12 Å with a force-based switching method^50^, and long-range electrostatics were computed using the particle-mesh Ewald method with a grid size of 0.16 Å and fourth-order interpolation^51^. Periodic boundary conditions were applied in all three directions. The equations of motion were integrated using the leapfrog algorithm with a 2 fs time step, saving trajectory data every 10 ps.

### Coarse-grained molecular dynamics (MD) simulations

Coarse-grained MD simulations were performed using Martini Maker in CHARMM-GUI, employing the same protein alignment and lipid composition as in the all-atom simulations^38,39,52,53^. Systems were solvated in water, with 150 mM NaCl added. The Martini3 force field was applied to describe the protein, lipids, water, and ions^54^. Coarse-grained MD simulation systems and system compositions are provided in Supplementary Table 1 and Supplementary Table 3, respectively.

Minimization and equilibration were carried out with gradually decreasing position restraints on the protein and lipid heads, employing the Berendsen barostat, v-rescale thermostat, and reaction-field electrostatics. Productive simulations of 100 µs were performed at 310.15 K in the NPT ensemble with a 20 fs time step, using Parrinello-Rahman pressure coupling. To preserve the protein secondary structure, an elastic network with a bond force constant of 500 kJ/(mol·nm^2^) was applied to the transmembrane helix backbones, with bond cutoffs set between 0.5 nm and 0.9 nm.

### Parallel cascade selection molecular dynamics (PaCS-MD)

PaCS-MD simulations were employed to study ion transport in mBOCT1^55^. Supplementary Table 2 lists all PaCS-MD systems used in this study. The initial conformations were selected from all-atom MD simulations, e.g. a frame with Fe^2+^ bound at the binding site. Starting from each selected conformation, multiple short MD simulations were performed for each system. MD frames were ranked based on the depth of Fe^2+^ in the lipid bilayer, measured by the distance between the ion and the center of mass of phosphate atoms in the extracellular lipid leaflet. The top-ranked frame was used as the starting point for subsequent simulation rounds. Each round consisted of 100 parallel MD simulations of 10 ns each, following the same settings as the all-atom simulations. This iterative process continued until Fe^2+^ was released to the intracellular side.

### Root mean square fluctuation (RMSF) and helicity analysis

The RMSF of the protein backbone was calculated using the gmx rmsf command in GROMACS to assess structural stability in the apo systems during all-atom MD simulations. Helicity of transmembrane residues was evaluated using the gmx helix command, with averages computed over the simulation duration.

### Root-mean-square deviation (RMSD)-based hierarchical clustering

Hierarchical clustering was used to identify representative structural conformations from both all-atom and coarse-grained MD simulations of apo systems. For all-atom simulations, trajectories were concatenated, and frames were extracted every 500 ps. For coarse-grained simulations, frames were extracted from 100 µs of trajectories at 20 ns intervals. All frames were superimposed using the backbone atoms/beads of TM3 (residues 127–145) and TM9 (residues 364–384). The pairwise RMSD matrix was computed for residues 220–302, encompassing TML6 and the intracellular domain.

Clustering was performed using SciPy’s hierarchical clustering algorithm, applying an RMSD cutoff of 10 Å for all-atom simulations and 20 Å for coarse-grained simulations^56^. The centroid frame of each cluster was selected, and coarse-grained models were converted back to all-atom models using CHARMM-GUI^38^.

### Hydrogen bond analysis

Hydrogen bonding dynamics were analyzed from concatenated PaCS-MD trajectories exported every 100 ps. Water molecules in the hydration shell of the Fe^2+^ ion were identified, and hydrogen bonds between the hydration shell and protein residues (D169 and E193) were calculated using the HBond plugin in VMD^57^. The criteria for hydrogen bonding were a donor-acceptor distance cutoff of 3.5 Å and a donor-H-acceptor angle cutoff of 150°.

## Supplementary Information

Supplementary Table 1. Non-biased molecular dynamics simulation systems.

Supplementary Table 2. All-atom parallel cascade selection molecular dynamics (PaCS-MD) simulation systems.

Supplementary Table 3. Molecular dynamics simulation system compositions.

Supplementary Table 4. Protein backbone root mean squared deviations (RMSD) from all-atom MD simulations.

Supplementary Table 5. Fe^2+^ ion RMSD from holo all-atom MD simulations.

Supplementary Table 6. mBOCT1–Fe^2+^ contact frequencies from MD simulations.

Supplementary Table 7. Penetration depth of Fe^2+^ ion in mBOCT1 from PaCS-MD.

Supplementary Figure 1. Sequence alignment between representative members of the SLC22 family.

Supplementary Figure 2. Sequence alignment of BOCT1 across various species.

Supplementary Figure 3. Secondary structure and transmembrane topology prediction from mBOCT1 sequence.

Supplementary Figure 4. Key amino acids that are vital for iron transport in mBOCT1.

Supplementary Figure 5. Different conformations of TML6 in MD simulations: the left panel shows the sideview, and the right panel shows the topview.

Supplementary Figure 6. Structural alignment of occluded mBOCT1 (cryo-EM, this study), outward-open mBOCT1 (MD simulations), and inward-open mBOCT1 (MD simulations).

Supplementary text: secondary structure and transmembrane topology prediction, simulation of apo mBOCT1, simulation of holo mBOCT1, parallel cascade selection molecular dynamics (PaCS-MD), homology modelling using hOCT1 as template.

Supplementary Movie 1: Fe^2+^ ion moving from the extracellular membrane lipid into the protein in model A, attracted by D336. The Fe^2+^ ion moves from D336 towards H200 in the last few seconds of the movie. D336 and H200 are labeled along the transport pathway.

In the movie, phosphate atoms in membrane lipids are represented as gray spheres, marking the membrane interface. Proteins are depicted as cyan cartoons, and Fe^2+^ ions are shown as red spheres.

## Data availability

The cryo-EM map has been deposited in the Electron Microscopy Data Bank (EMDB) under accession number EMD-60664, and the coordinates have been deposited in the Protein Data Bank (PDB) under accession number 9IL4. Source data are provided with this paper.

## Acknowledgments

We are grateful to Jiawei Wang from the School of Life Sciences, Bailong Xiao from the School of Pharmaceutical Sciences at Tsinghua University, and Daohua Jiang from the University of Chinese Academy of Science for their helpful discussions and sharing of reagents. We also acknowledge Shuimu BioSciences for their cooperation in cryo-EM data collection. This work has been supported by the National Natural Science Foundation of China (Grants 32130048, 92157301 to L.C.), the Ministry of Science and Technology of China National Key R&D Programs (Grant 2022YFA0806503 and 2024YFA1306103 to L.C.), a private donation from the Tsinghua University Education Foundation (Grant 202011 to L.C.), a Collaborative Fund from CytoCraft Biotech Co. and the Tsinghua-Foshan Innovation Special Fund (Grant 20229990133 to L.C.), as well as the Tsinghua Precision Medicine Foundation (Grant No.2022TS013). W.W. was partially supported by the SGUnited Jobs Initiative (Grant P20J3d1006) from the National Research Foundation Singapore. Both W.W. and H.F. were supported by the Biomedical Research Council of A*STAR, Singapore. The computational work for this article was performed using the resources of the National Supercomputing Centre, Singapore (https://www.nscc.sg).

## Author contributions

L.C. conceived the project. J.C., L.C., X.L., and H.F. designed the experiments. J.C. performed the purification experiments. J.C. and S.Z. prepared the cryo-EM samples. J.C. collected the cryo-EM data. S.Z. and J.C. calculated the EM map. J.C., X.L., and S.Z. built the model. J.C. carried out the functional assays *in vitro* and *in vivo*. W.W. and H.F. conducted the MD simulations. J.C., S.Z., W.W., L.T., L.T., K.G., B.L., A.E., H.F., X.L., and L.C. analyzed and discussed the data. L.C., J.C., X.L., and H.F. wrote the manuscript.

## Supplementary information

**Extended Data Fig. 1.**
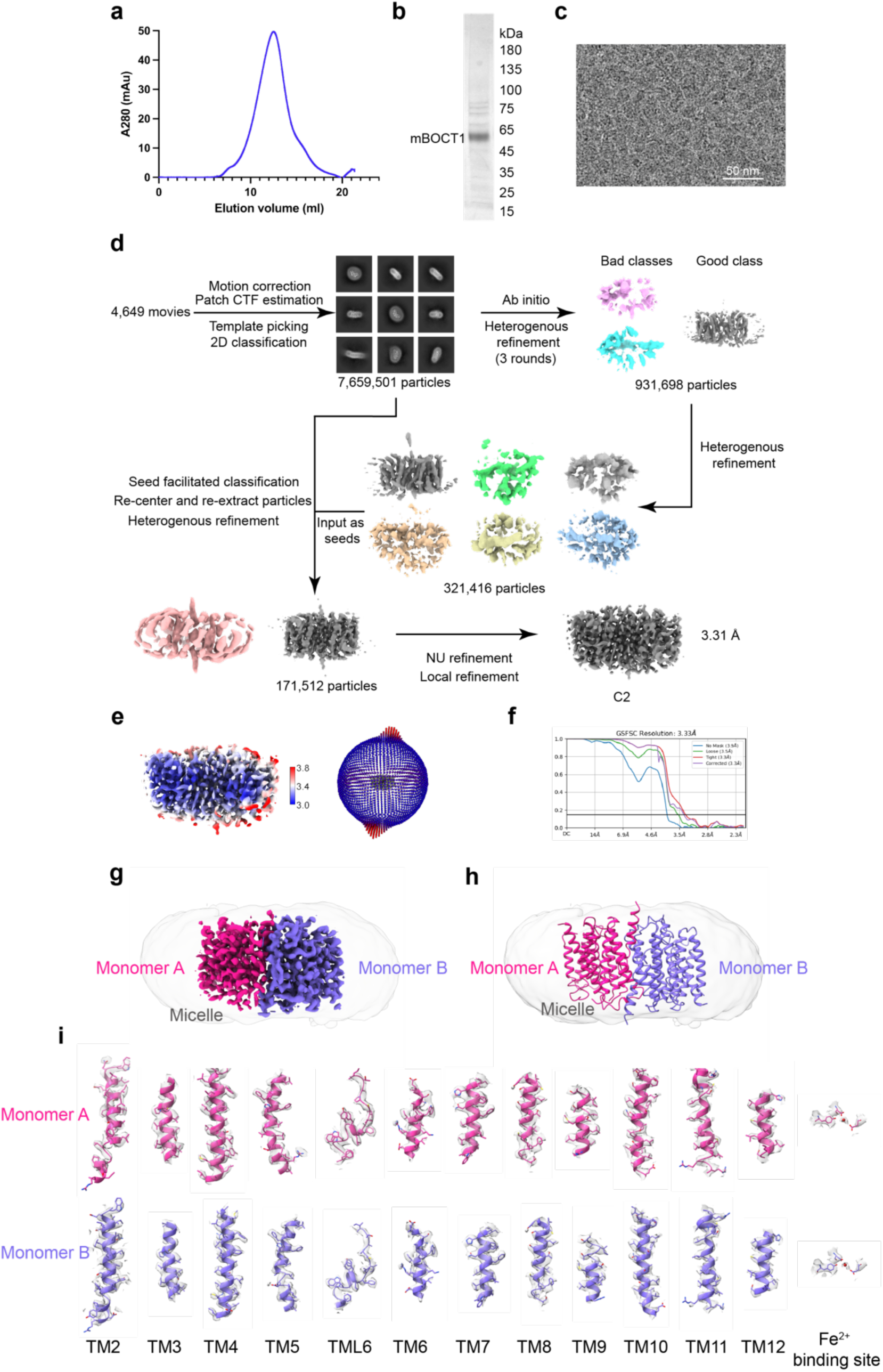
Cryo-EM structure determination of mBOCT1. **a**, Size exclusion chromatography (SEC) profile of mBOCT1. **b**, SDS-PAGE analysis of peak fractions from the SEC of mBOCT1. **c**, Representative cryo-EM image of mBOCT1 captured at 300 kV. **d**, Workflow for single-particle analysis of mBOCT1 using cryoSPARC. **e**, Local resolution estimation and angular distribution of mBOCT1 in cryo-EM data. **f**, FSC curve of mBOCT1. **g**, Cryo-EM map of the mBOCT1 dimer. **h**, Model of the mBOCT1 dimer. **i**, Density map of TMs and Fe^2+^ binding site in mBOCT1, depicted as a gray surface.

**Extended Data Fig. 2.**
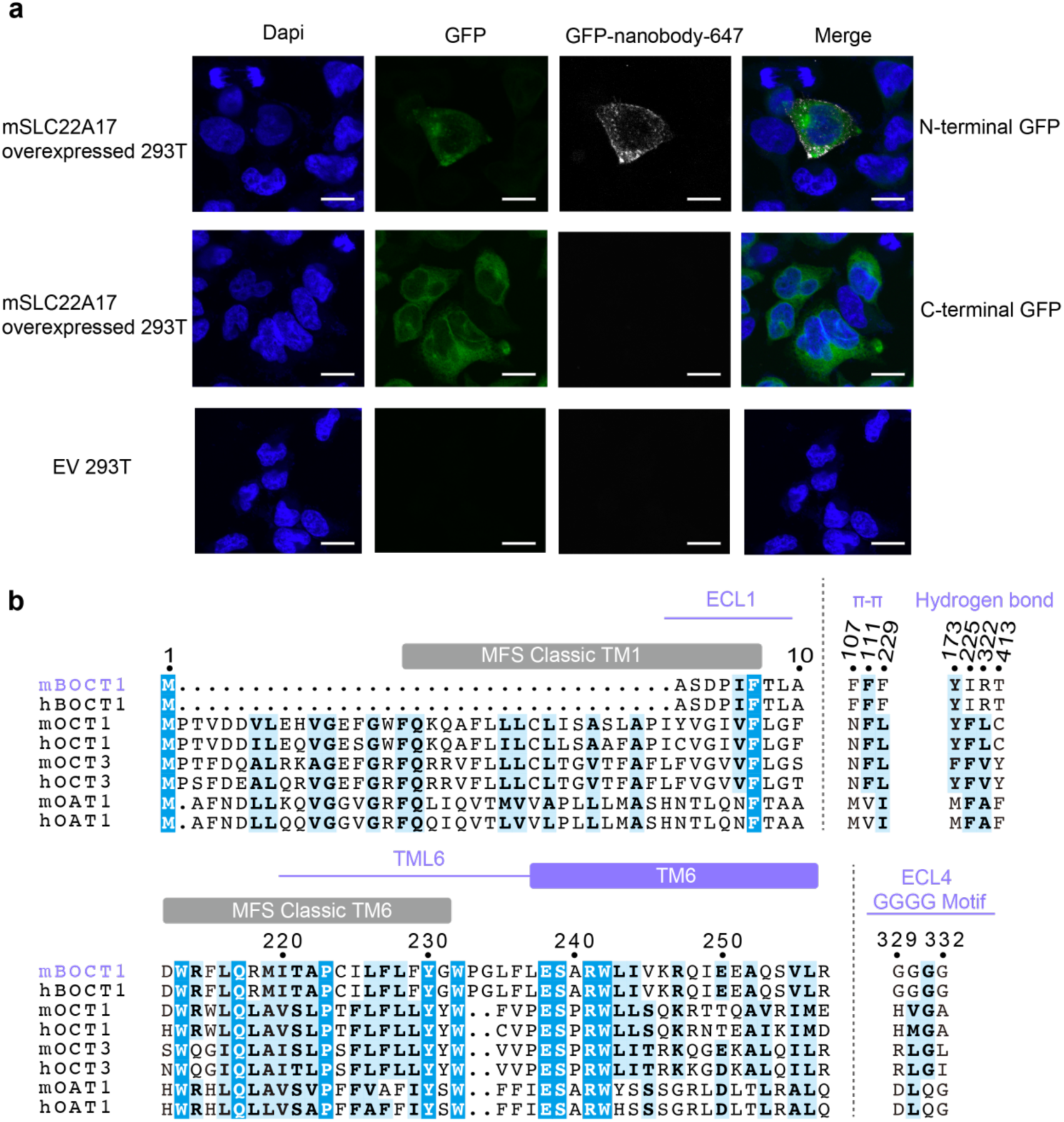
Structure verification *in vivo* and sequence alignment of mBOCT1. **a**, Surface staining of mBOCT1 tagged with N-terminal or C-terminal GFP in stable 293T cell lines and EV 293T cell lines via immunofluorescence. DAPI indicates cell nuclei, GFP represents mBOCT1-GFP, and GFP-nanobody-647 shows the GFP nanobody labeled with Alexa Fluor 647. Scale bar: 10 µm. **b**, Sequence alignment of unique regions in mBOCT1 compared to selected SLC22 family members. TMs are represented as rectangles above the sequences, loops as lines. Secondary structural features of mBOCT1 are shown in purple, while those of traditional MFS transporters are in gray.

**Extended Data Fig. 3.**
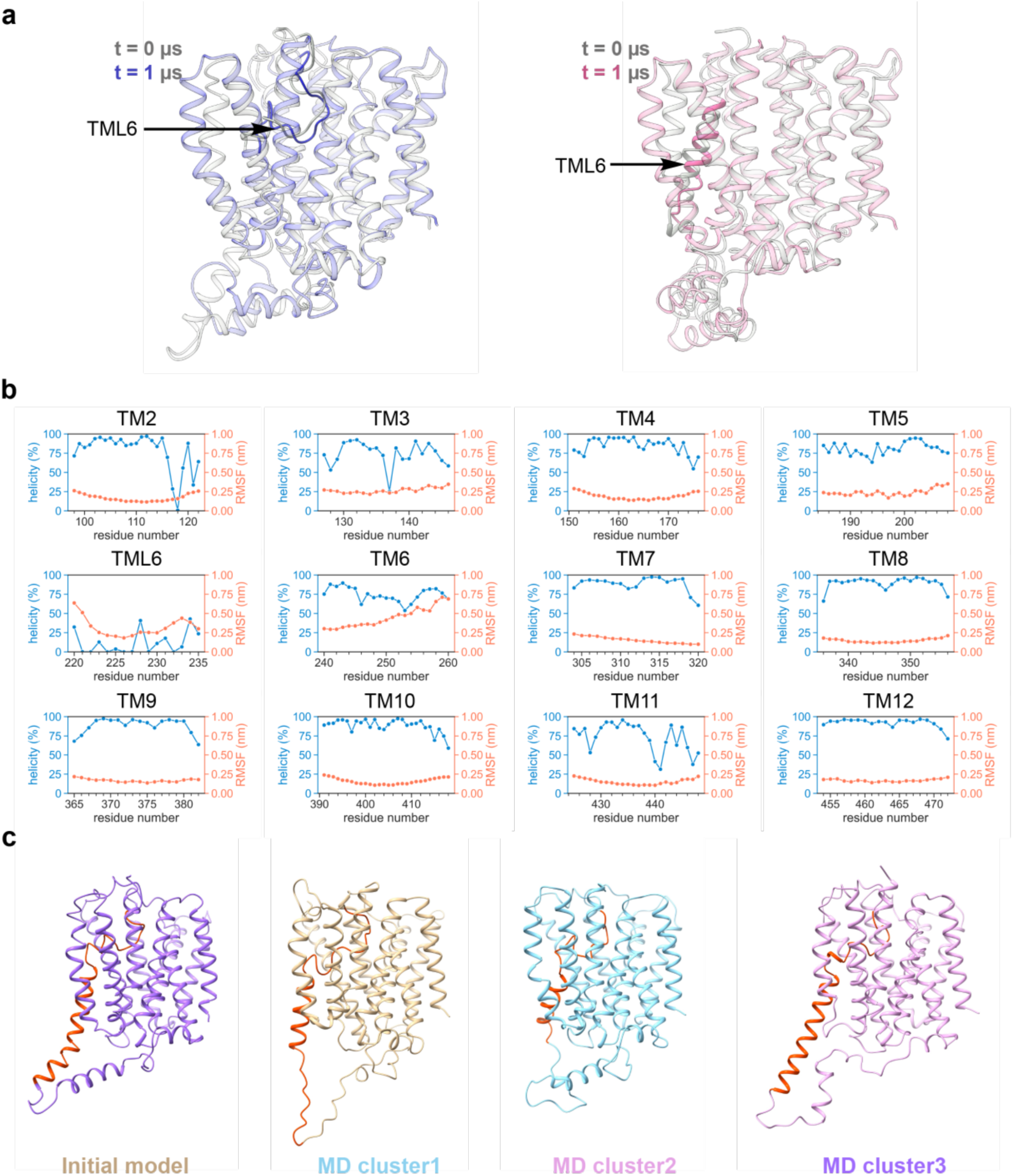
Structural stability of TML6 and TM6 in MD simulations. a, Stability of TML6 in loop or helix form observed in apo MD simulations (System 1 and 2 in Supplementary Table 1). The time scale is 1 µs. TML6 is marked by an arrow, and other structural segments are set to 80% transparency. **b**, RMSF and helicity of the 11 TMs in mBOCT1 from all-atom MD simulations. **c**, Various conformational states of TML6 and TM6 in mBOCT1 observed in MD simulations. TML6 and TM6 are highlighted in red.

**Extended Data Fig. 4.**
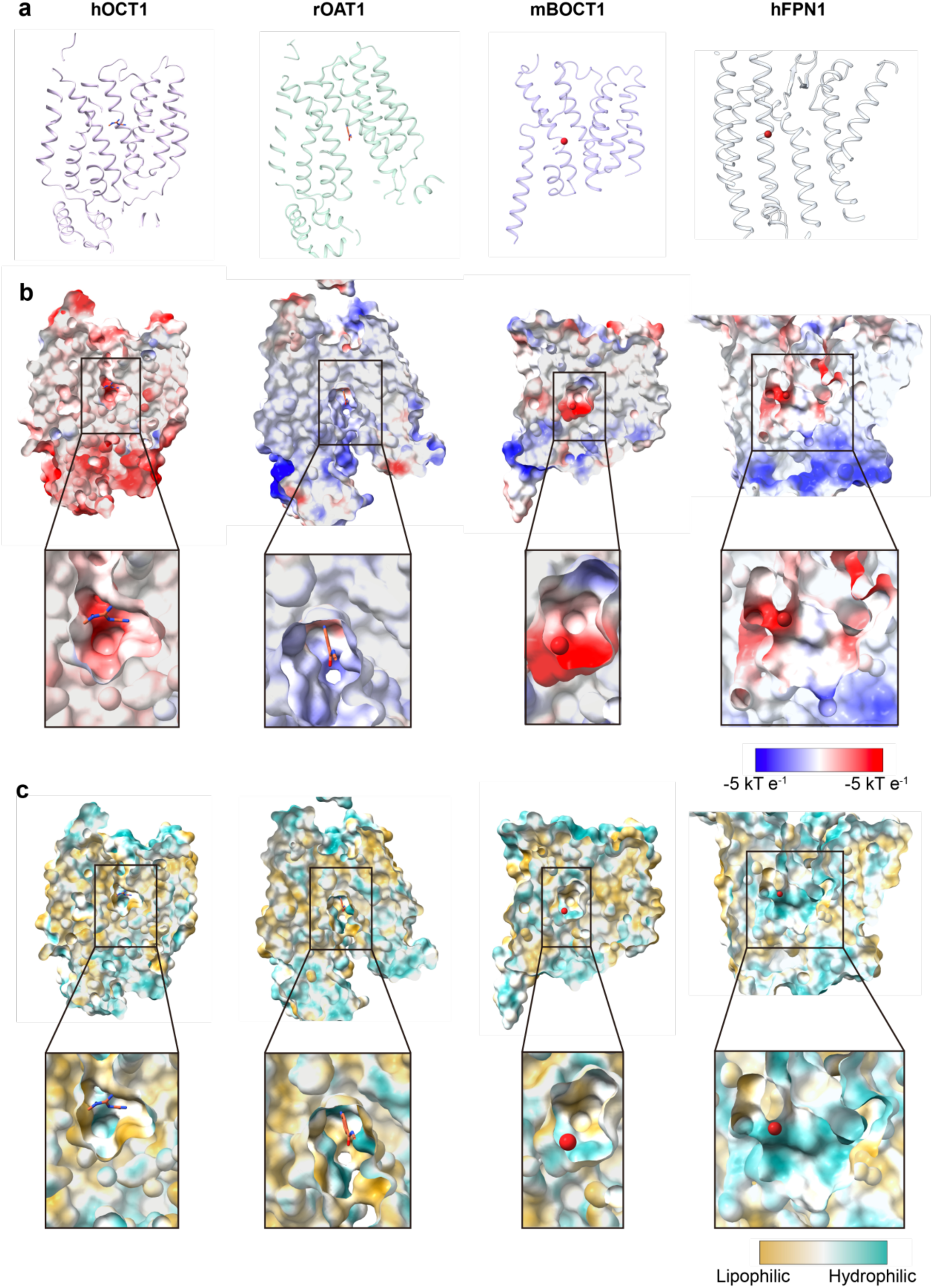
Structural comparison of mBOCT1 with selected SLC22 members and hFPN1. **a**, Models of hOCT1 (PDB: 8JTT), rOAT1 (PDB: 8SDY), mBOCT1 (this study), and hFPN1 (PDB: 6WBV). **b**, Corresponding sliced views of surface potential and magnified views of the binding pocket. A color key for electrostatic potential is displayed below. **c**, Sliced views of hydrophilicity analysis and magnified views of the binding pocket. A color key for hydrophobicity is displayed below.

**Extended Data Fig. 5.**
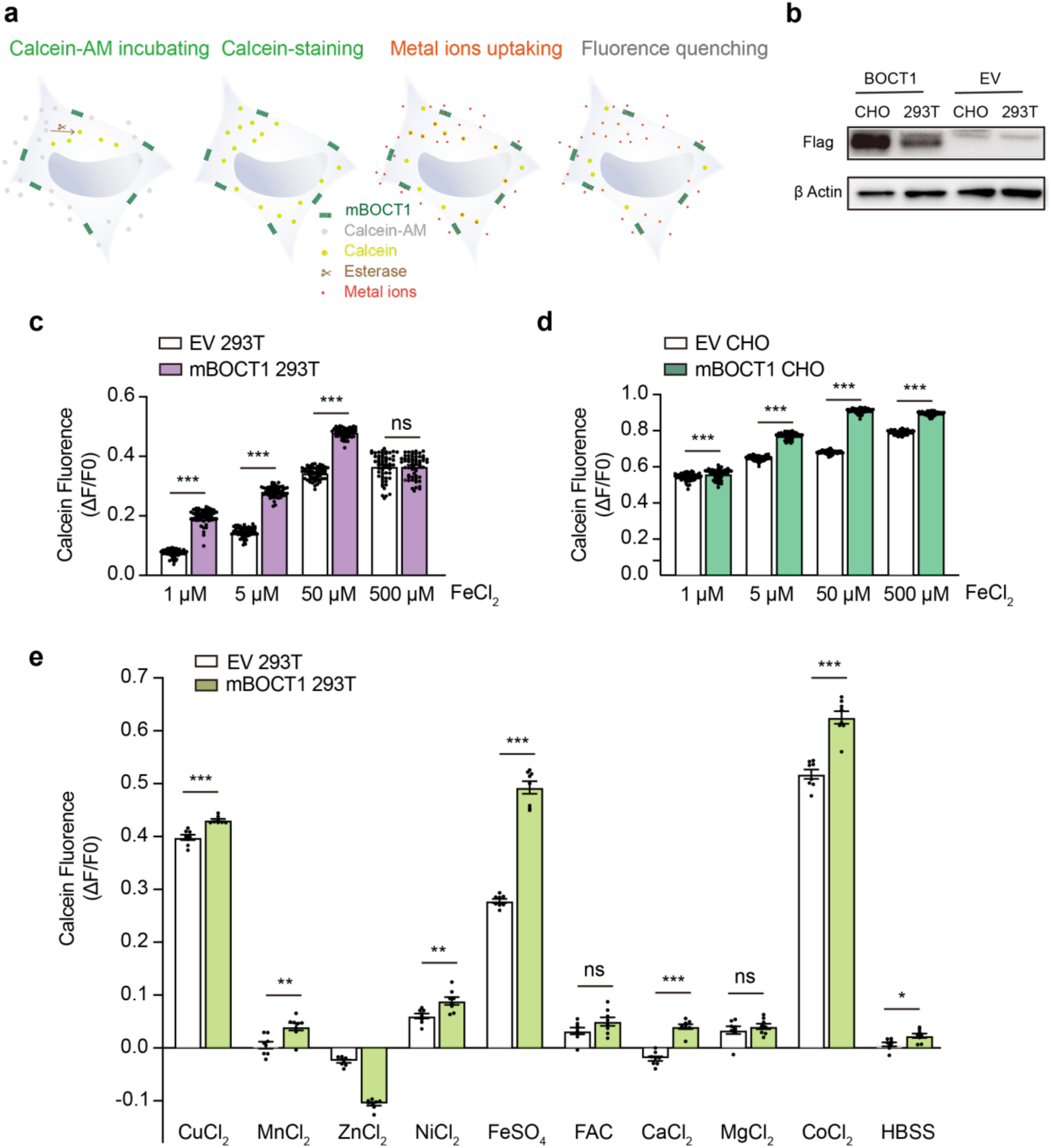
Cellular uptake of mBOCT1 with metal ions compounds. a, Schematic of the cellular uptake assay using calcein. Non-fluorescent calcein-AM was incubated with mBOCT1 293T stable cell lines and converted into fluorescent calcein by esterases. Fluorescence quenching occurred upon metal ion transport into cells. **b**, Western blot validation of stable mBOCT1 overexpression in 293T and CHO cells. mBOCT1 was fused with an N-terminal Flag tag. **c**, Cellular uptake assay of mBOCT1-expressing HEK 293T stable cell lines with FeCl_2_. Data are shown as Mean ± SEM from n > 3 independent experiments. **d**, Cellular uptake assay of mBOCT1-expressing CHO stable cell lines with FeCl_2_. Data are shown as Mean ± SEM from n > 3 independent experiments. **e**, Cellular uptake assay of HEK 293T stable cell lines with various metal ion compounds. Data are shown as Mean ± SEM from n > 3 independent experiments. Significance for panels c–e was analyzed using an unpaired *t*-test. Significance levels: \**p* < 0.05, \*\**p* < 0.01, and \*\*\**p* < 0.001.

**Extended Data Fig. 6.**
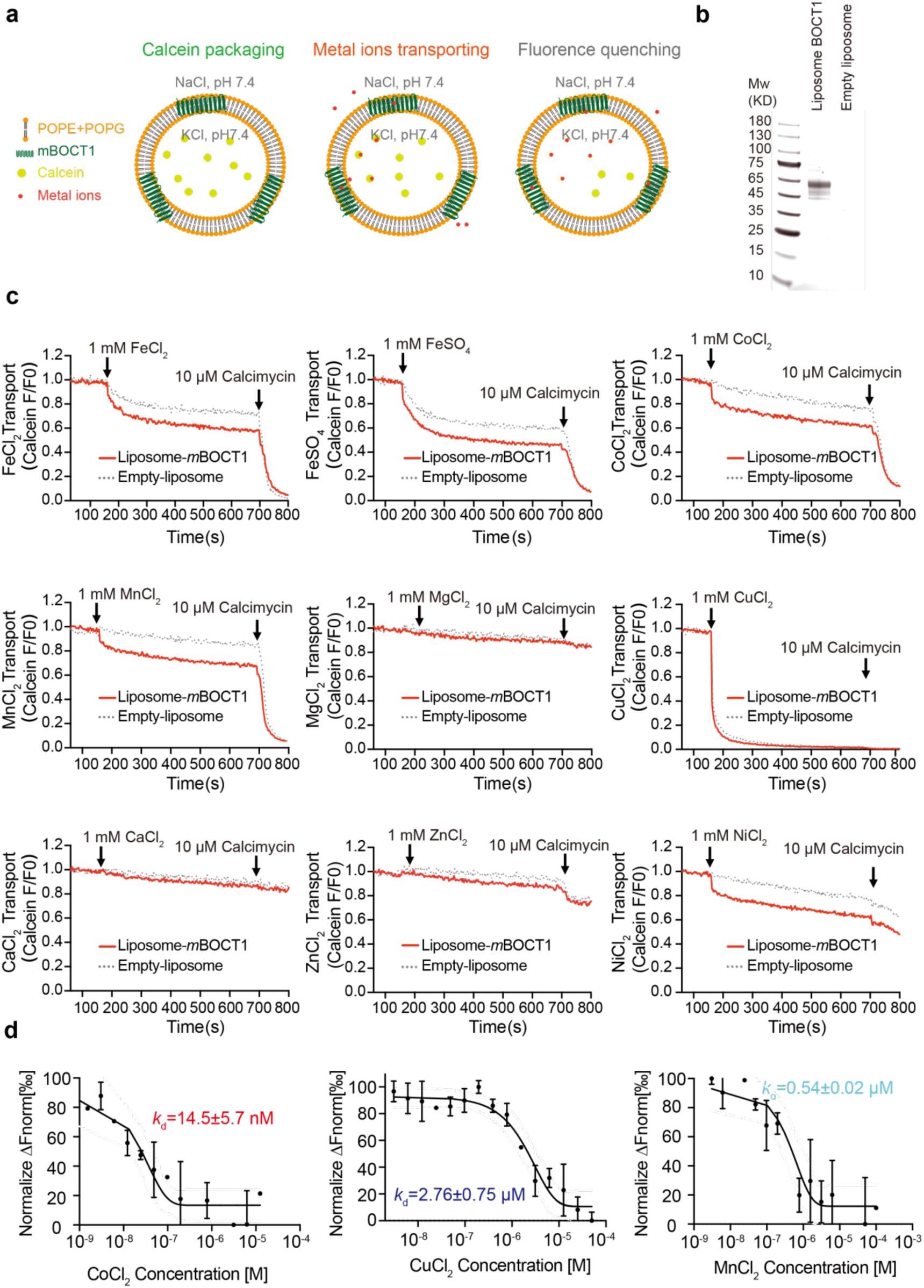
Liposome transport and MST binding assay of mBOCT1 with metal ions compounds. **a**, Schematic of the liposome transport assay using calcein fluorescence as an indicator. Purified mBOCT1 was reconstituted into liposomes composed of POPE and POPG, containing calcein. Fluorescence quenching occurs upon metal ion transport or when the ionophore calcimycin is added. **b**, Verification of mBOCT1 reconstitution by SDS-PAGE. **c**, Representative dynamic transport curves of various metal ions in the mBOCT1 liposome system. Data were obtained from three independent replicates. **d**, Binding curves for different metal ions with mBOCT1 measured by MST. Data were obtained from three independent replicates.

**Extended Data Fig. 7.**
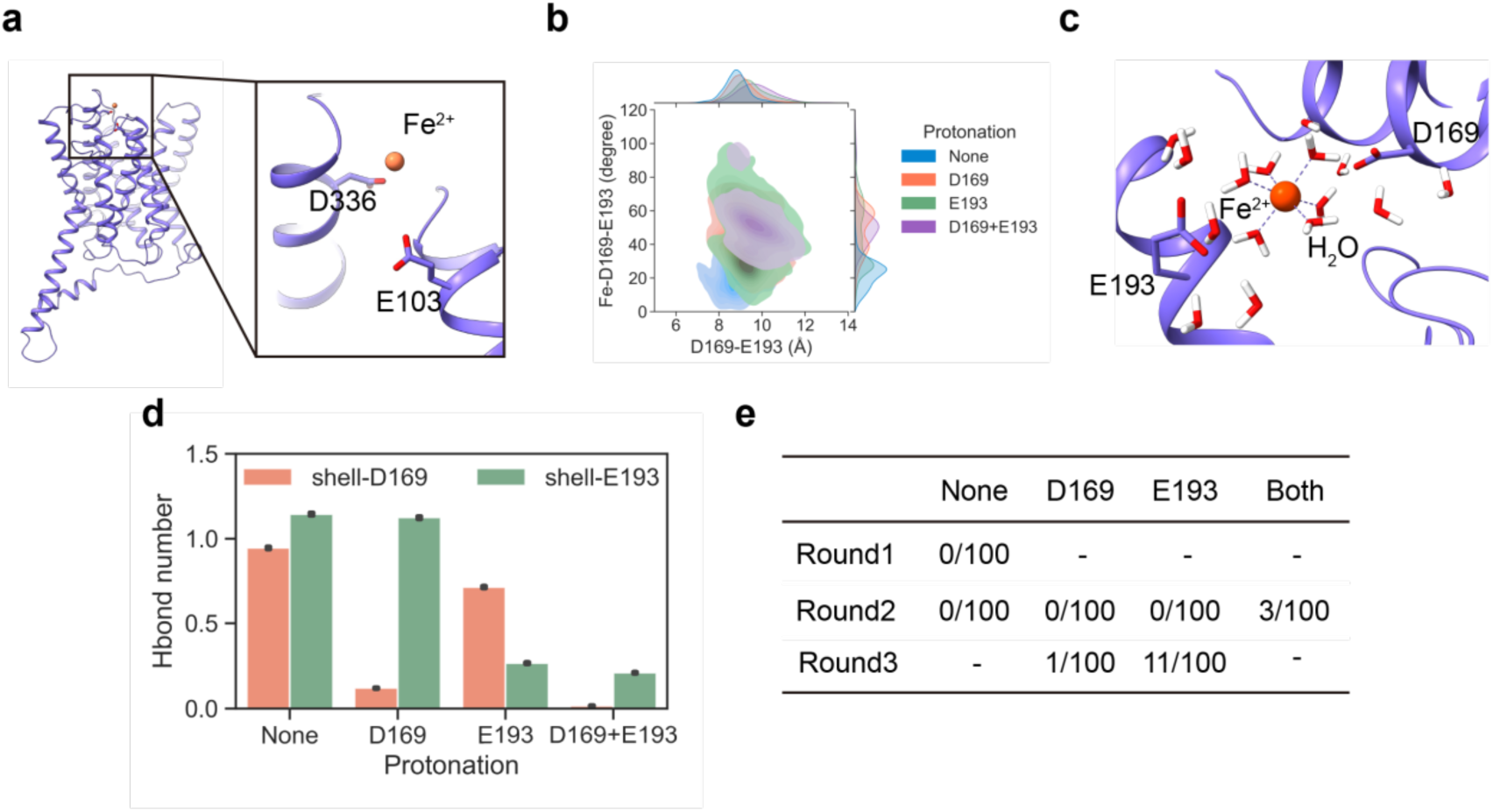
Effect of D336 and the protonation of D169 and E193 on iron transport as indicated by MD simulations. **a**, Residues at the entrance of the mBOCT1 transport pathway when Fe^2+^ enters the transporter in MD simulations. **b**, Distance and angle changes between D169-E193 and Fe^2+^-D169-E193 upon protonation of D169 or E193. **c**, Surrounding water molecules around the Fe^2+^ binding site in the mBOCT1 transport pathway. **d**, Hydrogen bonds between the hydration shell and D169/E193 upon protonation of D169 or E193. **e**, Number of Fe^2+^ release events (out of 100 parallel runs) in three rounds of PaCS-MD simulations upon protonation of D169 or E193.

**Extended Data Fig. 8.**
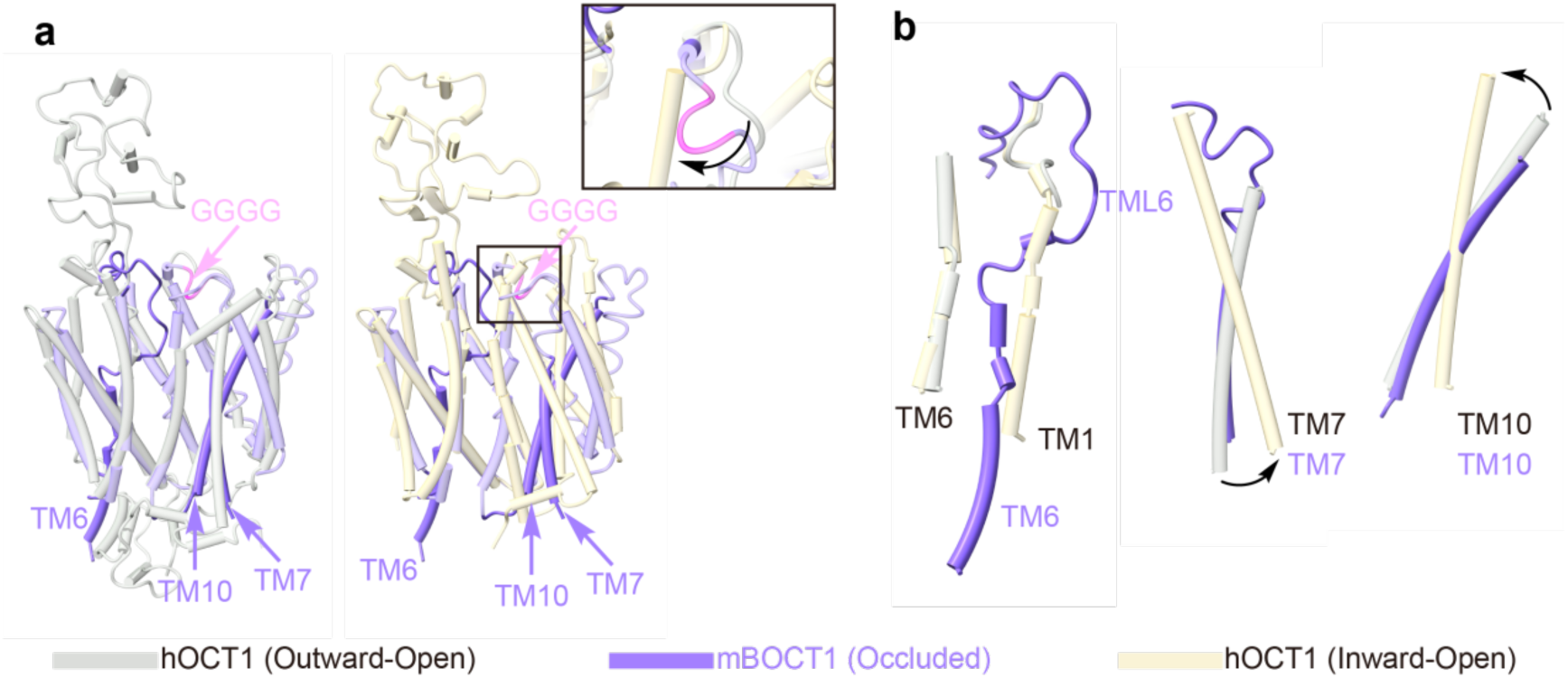
Structure comparisons of mBOCT1 with hOCT1 in different conformations. **a**, Structural alignments of mBOCT1 with outward-open hOCT1 (PDB: 8et6) and inward-open hOCT1 (PDB: 8sc1). Important helices and motifs essential for mBOCT1 transport are highlighted. **b**, Structural alignments of TML6, TM6, TM7, and TM10 of mBOCT1 with the corresponding TMs in outward-open and inward-open hOCT1.

